# Structural analysis of chlorogenic acid from red clover (*Trifolium pratense*) extract

**DOI:** 10.64898/2026.05.21.726747

**Authors:** Fedorova Anastasia Mikhailovna, Milentyeva Irina Sergeevna, Asyakina Lyudmila Konstantinovna, Prosekov Alexander Yuryevich

## Abstract

This paper presents the results of a structural analysis of chlorogenic acid isolated from a 70% ethanol extract of red clover (*Trifolium pratense*) callus culture. X-ray phase analysis showed that the sample was crystalline and single-phase and crystallized in an orthorhombic unit cell with the following parameters: a = 36.7548(5) Å, b = 11.0770(3) Å, c = 7.7947(2) Å, V = 3173.46(11) Å^3^, R-Bragg = 0.347 %, R_exp_ = 4.75 %, R_wp_ = 5.83 %, R_p_ = 4.39 %, GOF = 1.23 %. NMR spectroscopy data (^1^H, ^13^C{^1^H}, 2D ^1^H^1^H-COSY, ^1^H^13^C-HSQC, ^1^H^13^C-HMBC) confirmed that the chemical structure and purity of the sample fully corresponded to chlorogenic acid, with no chemical impurities detected. Complete proton and carbon atom assignments are provided.

## Introduction

There is currently a steady increase in interest in biologically active plant-derived compounds as a basis for developing modern pharmaceuticals and nutraceuticals [1]. Natural compounds are characterized by a broad spectrum of therapeutic activities and, as a rule, low toxicity, making them promising candidates for pharmaceutical development. Among the diverse class of polyphenolic compounds, chlorogenic acid (a caffeic and quinic acid ester) occupies a special place because of its wide range of bioactivities, including antioxidant [2], anti-inflammatory [3], hepatoprotective [4], and hypoglycemic properties [5]. Interest in this compound is also associated with its ability to modulate the activity of enzymes involved in carbohydrate metabolism and to exert neuroprotective effects [6].

A promising source of this compound is red clover (*Trifolium pratense*), a perennial herbaceous plant of the legume family traditionally used in folk medicine because of its high content of isoflavonoids, phenolic acids, and other valuable phytonutrients [7]. Of particular interest are biotechnologically derived callus, suspension, and root cultures *in vitro*, which make it possible to obtain standardized plant material with a controlled content of target metabolites, regardless of seasonal, climatic, and environmental factors [8, 9]. At earlier stages of the present study, an extract of red clover callus culture was obtained, from which chlorogenic acid with a purity of at least 95% was isolated [10].

However, the efficacy and safety of individual compounds are largely determined by their structural features. For a comprehensive characterization of polyphenols, including chlorogenic acid, it is critically important to establish not only their molecular structure but also their three-dimensional structure structure, since crystal packing and molecular conformation can affect reactivity and bioavailability [11]. Modern physicochemical methods, such as nuclear magnetic resonance (NMR) spectroscopy and X-ray diffraction analysis, offer unique opportunities for detailed structural analysis [12]. NMR spectroscopy enables highly accurate confirmation of the chemical structure and purity of a compound in solution, whereas powder X-ray diffraction provides information on the phase composition and crystal lattice parameters of a solid sample [13]. The combined use of these complementary approaches provides the most complete understanding of the substance’s structure.

The aim of the present study was to investigate the molecular and spatial structure of chlorogenic acid isolated from a 60% ethanol extract of red clover (*Trifolium pratense*) callus culture using NMR spectroscopy (^1^H, ^13^C{^1^H}, 2D ^1^H^1^H-COSY, ^1^H^13^C-HSQC, ^1^H^13^C-HMBC) and X-ray diffraction analysis. To achieve this aim, the following objectives were addressed: to determine the phase composition and crystal structure of the sample by powder X-ray diffraction; to establish the molecular structure and confirm the purity of chlorogenic acid using one- and two-dimensional NMR spectroscopy; and to perform complete assignment of the proton and carbon signals in the ^1^H and ^13^C{^1^H} NMR spectra.

## Materials and Methods

The object of the study was chlorogenic acid isolated from a 70% ethanol extract of red clover (*Trifolium pratense*) callus culture, with a purity of at least 95%.

The red clover (*Trifolium pratense*) callus culture and extract were obtained at earlier stages of the study. The procedures used to obtain the callus culture and extract are described by L. S. Dyshlyuk et al. [14]. Calluses were obtained from aseptically damaged seedlings grown from seeds acquired from the collection of E. K. Sirotkin, Russia. The seedlings were cultivated on solidified Gamborg medium supplemented with kinetin (2.00 mg/L), 6-BAP (0.10 mg/L), IAA (2.00 mg/L), and 2,4-D (2.00 mg/L) [15]. Extracts from the callus cultures were prepared by 6 h extraction under reflux condenser in a water bath at 70.00 ± 5.00°C, using a 70.00% aqueous ethanol solution as the extractant. The isolation of chlorogenic acid from the red clover (*Trifolium pratense*) callus culture extract is described by E. R. Faskhutdinova et al. [10]. The purity of chlorogenic acid and biochanin A was at least 95.00%.

Crystal structure analysis was performed using a Bruker D8 Advance X-ray diffractometer with CuKα radiation and a LynxEye position-sensitive detector in reflection geometry with sample rotation. Data were collected using the Bruker DIFFRACplus software package and analyzed with EVA and TOPAS V5.0.

For NMR spectroscopy, 20 mg portions of the study samples – rutin, quercetin, baicalin, trans-cinnamic acid, and chlorogenic acid – were dissolved in 600 μL of dimethyl sulfoxide (DMSO) at 25°C. The resulting solutions were placed in a Bruker Avance III HD 500 NMR spectrometer for spectra acquisition. The operating frequencies of the NMR spectrometer are shown in Table 1.

**Table 1.**
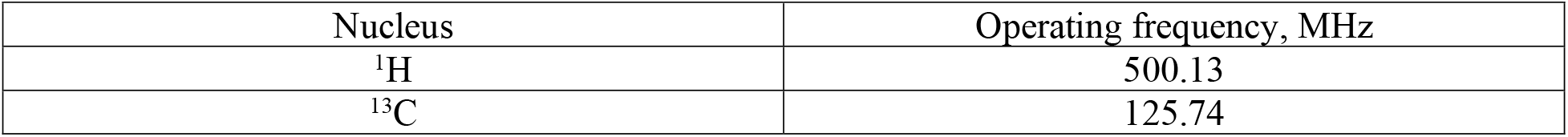
Operating frequencies used in the study.

The ^13^C{^1^H} spectra were recorded with suppression of spin–spin interactions with ^1^H nuclei, using ^1^H decoupling and phase selection (J-Modulated Spin-Echo) to distinguish the signals of CH, CH_3_иCH_2_, and C groups in the spectra. For signal assignment in the ^1^H and ^13^C{^1^H} spectra, additional NMR spectra were acquired: 2D ^1^H^1^H-COSY, ^1^H^13^C-HSQC, and ^1^H^13^C-HMBC.

### Results and Discussion

The analysis showed that the chlorogenic acid sample was crystalline and single-phase (Figure 1, Table 2).

**Table 2.** Intensity and position of selected peaks with a relative intensity greater than 5% for the chlorogenic acid sample.

| Pos. [2Th] | Height [cts] | FWHM Left [2Th] | d-spacing [Å] | Rel. Int. [%] |
| --- | --- | --- | --- | --- |
| 4.799 | 3284.59 | 0.100 | 18.40 | 72.6 |
| 9.622 | 2039.36 | 0.100 | 9.18 | 43.6 |
| 14.461 | 605.65 | 0.136 | 6.12 | 9.8 |
| 14.706 | 624.63 | 0.138 | 6.02 | 10.2 |
| 15.681 | 472.58 | 0.100 | 5.65 | 6.4 |
| 15.998 | 2645.73 | 0.100 | 5.54 | 56.5 |
| 16.179 | 1355.75 | 0.100 | 5.47 | 26.5 |
| 16.717 | 948.58 | 0.100 | 5.30 | 16.9 |
| 17.568 | 1987.21 | 0.100 | 5.04 | 40.9 |
| 18.420 | 1091.65 | 0.153 | 4.81 | 19.9 |
| 18.699 | 1319.39 | 0.147 | 4.74 | 25.1 |
| 19.305 | 622.36 | 0.100 | 4.59 | 8.7 |
| 19.809 | 1072.76 | 0.154 | 4.48 | 19.1 |
| 20.096 | 1138.92 | 0.156 | 4.41 | 20.6 |
| 20.383 | 510.07 | 0.158 | 4.35 | 6.1 |
| 20.927 | 3032.50 | 0.100 | 4.24 | 64.5 |
| 21.926 | 980.31 | 0.100 | 4.05 | 17.4 |
| 23.204 | 759.24 | 0.116 | 3.83 | 11.9 |
| 23.864 | 688.11 | 0.100 | 3.73 | 9.8 |
| 24.318 | 1025.44 | 0.100 | 3.66 | 17.4 |
| 24.684 | 2582.02 | 0.100 | 3.60 | 53.2 |
| 25.285 | 1705.68 | 0.165 | 3.52 | 32.8 |
| 25.521 | 4610.79 | 0.100 | 3.49 | 100.0 |
| 25.896 | 582.27 | 0.100 | 3.44 | 6.8 |
| 26.092 | 1335.13 | 0.100 | 3.41 | 24.2 |
| 26.812 | 515.96 | 0.100 | 3.32 | 5.4 |
| 27.150 | 512.48 | 0.178 | 3.28 | 5.3 |
| 29.475 | 576.93 | 0.101 | 3.03 | 6.3 |
| 30.048 | 951.17 | 0.160 | 2.97 | 14.9 |
| 30.286 | 2900.35 | 0.100 | 2.95 | 60.0 |
| 31.374 | 593.03 | 0.100 | 2.85 | 6.8 |
| 31.754 | 734.38 | 0.176 | 2.82 | 10.2 |
| 32.406 | 536.46 | 0.172 | 2.76 | 5.5 |
| 34.694 | 726.45 | 0.100 | 2.58 | 8.3 |

**Figure 1.**
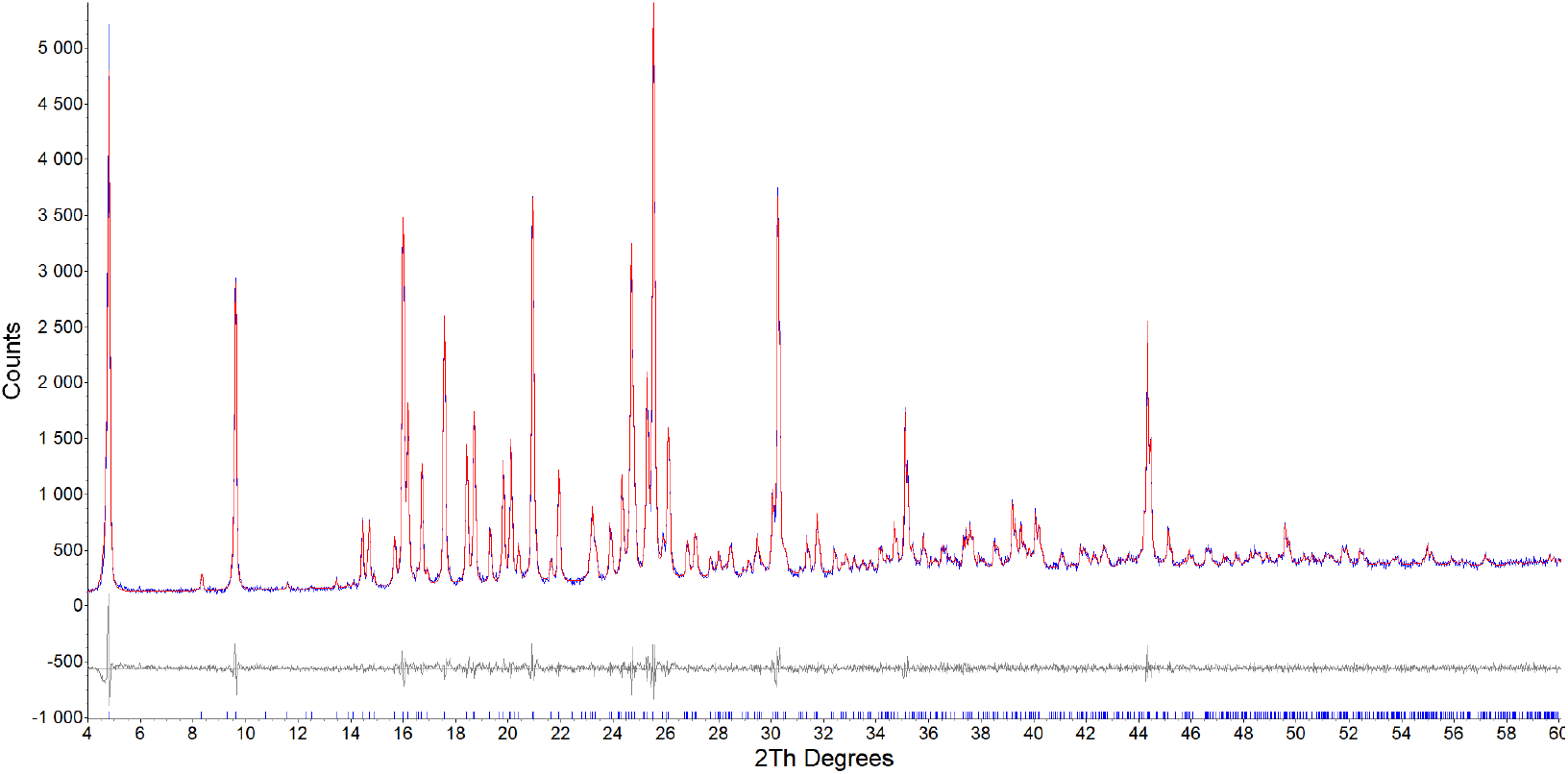
Experimental diffractogram of the chlorogenic acid sample (blue curve), theoretical diffractogram calculated for the corresponding unit cell (red line), and the difference curve between them (gray line). Vertical tick marks indicate the calculated peak positions.

Indexing results showed that the chlorogenic acid sample crystallized in an orthorhombic unit cell with the following parameters: a = 36.7548(5) Å, b = 11.0770(3) Å, c = 7.7947(2) Å, V = 3173.46(11) Å^3^, R-Bragg = 0.347 %, R_exp_ = 4.75 %, R_wp_ = 5.83 %, R_p_ = 4.39 %, GOF = 1.23 %.

The diffraction analysis showed that the chlorogenic acid sample under study corresponded to the stated compound, chlorogenic acid.

Additional structural studies of the chlorogenic acid sample were then performed using NMR spectroscopy.

Figure 3 shows the NMR spectra of the chlorogenic acid sample, which was presumed to have the structure of chlorogenic acid: ^1^H, ^13^C{^1^H}, 2D ^1^H^1^H-COSY, ^1^H^13^C-HSQC, ^1^H^13^C-HMBC.

The results presented above (Figure 3) show that the ^1^H, 13C{^1^H}, ^1^H^1^H-COSY, ^1^H^13^C-HSQC, and ^1^H^13^C-HMBC NMR spectra of the sample correspond to the target compound, chlorogenic acid. No chemical impurities were observed. In terms of the number of signals, their spectral positions (chemical shifts), integral intensities and their ratios, signal multiplicity, and spin– spin coupling constants (SSCCs), the obtained spectra are consistent with the target compound (Figure 4).

**Figure 3.**
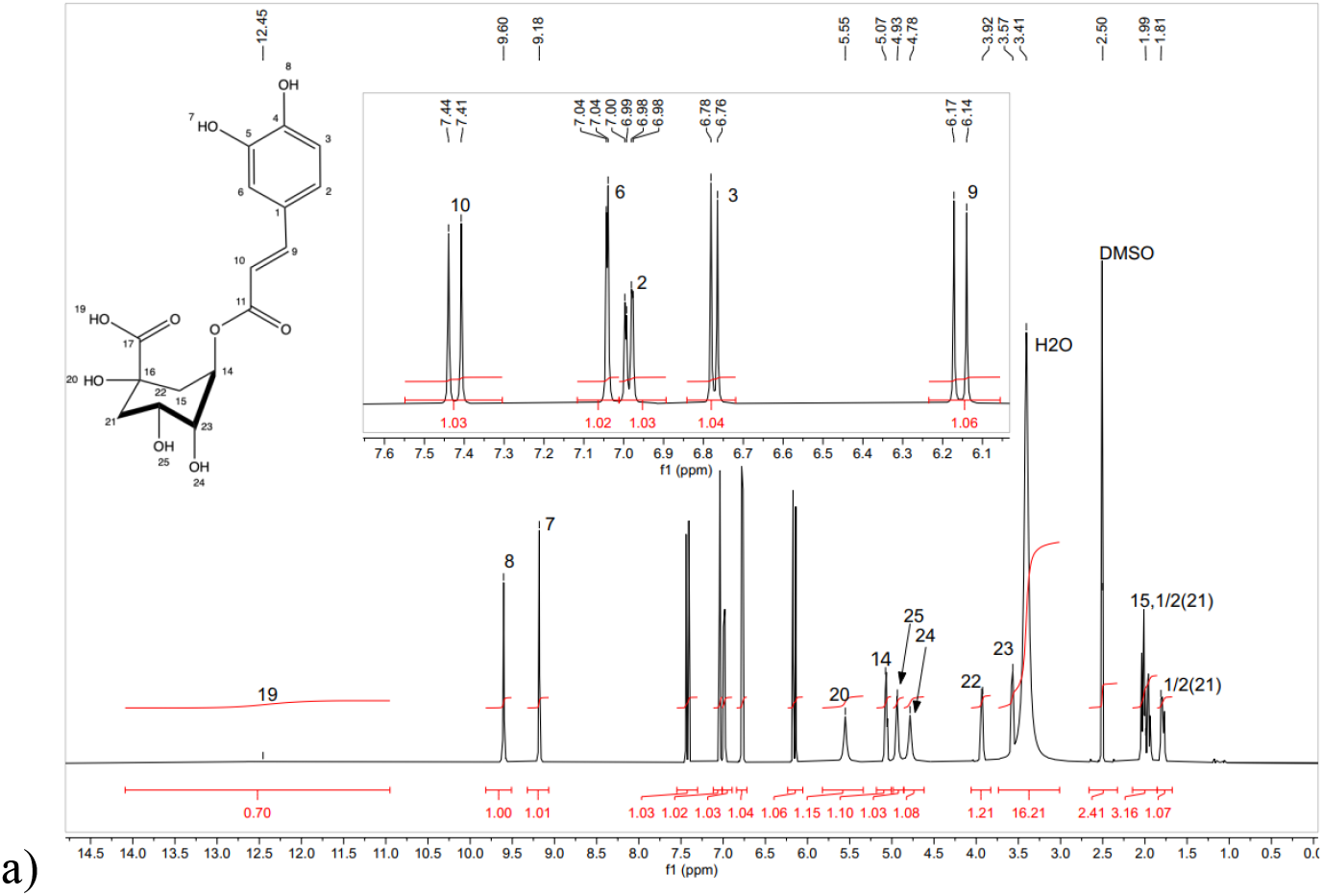
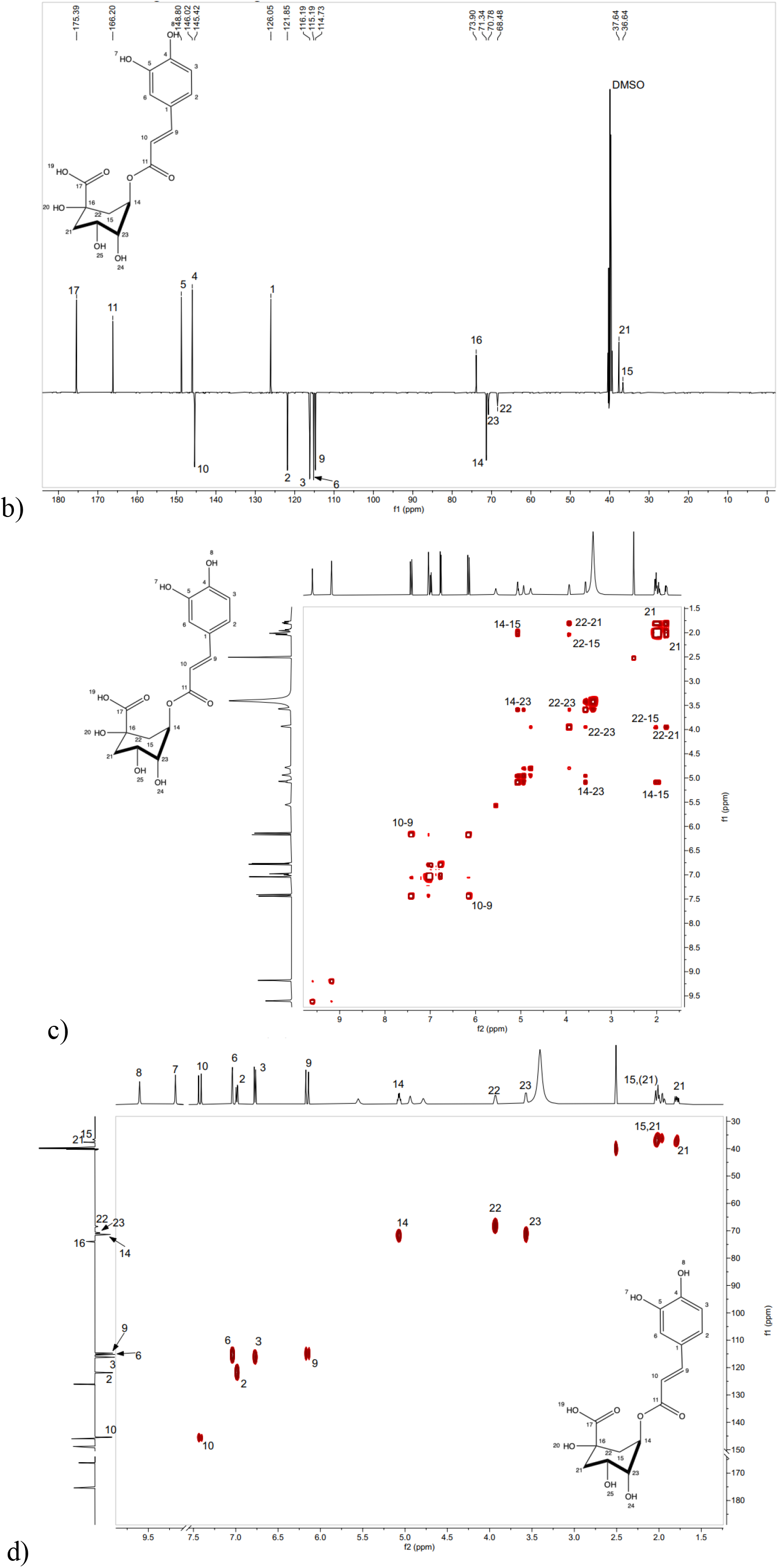
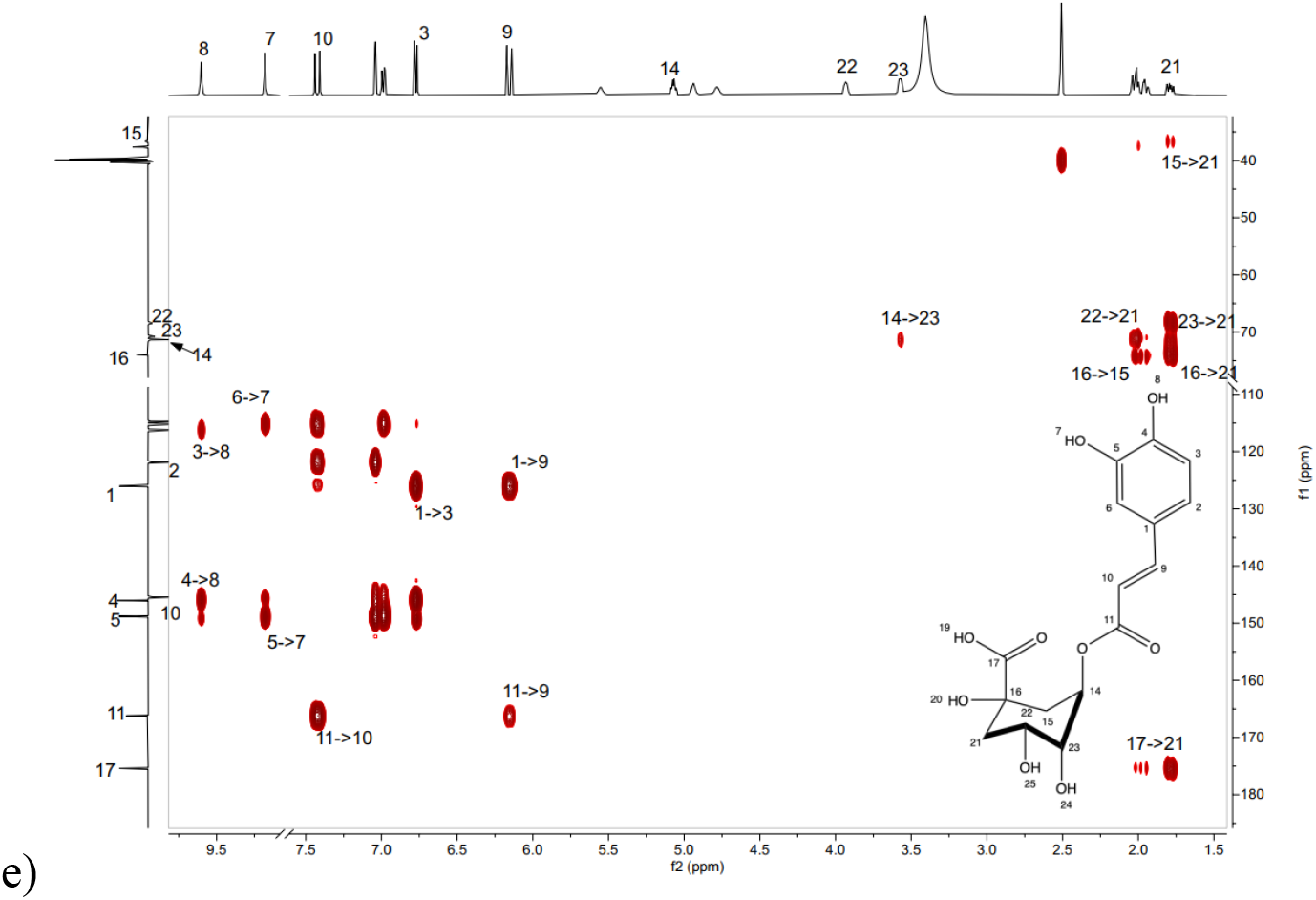
NMR spectra of the chlorogenic acid sample: a – ^1^H, b – ^13^C{^1^H}, c – 2D ^1^H^1^H-COSY, d – ^1^H^13^C-HSQC, e – ^1^H^13^C-HMBC

**Figure 4.**
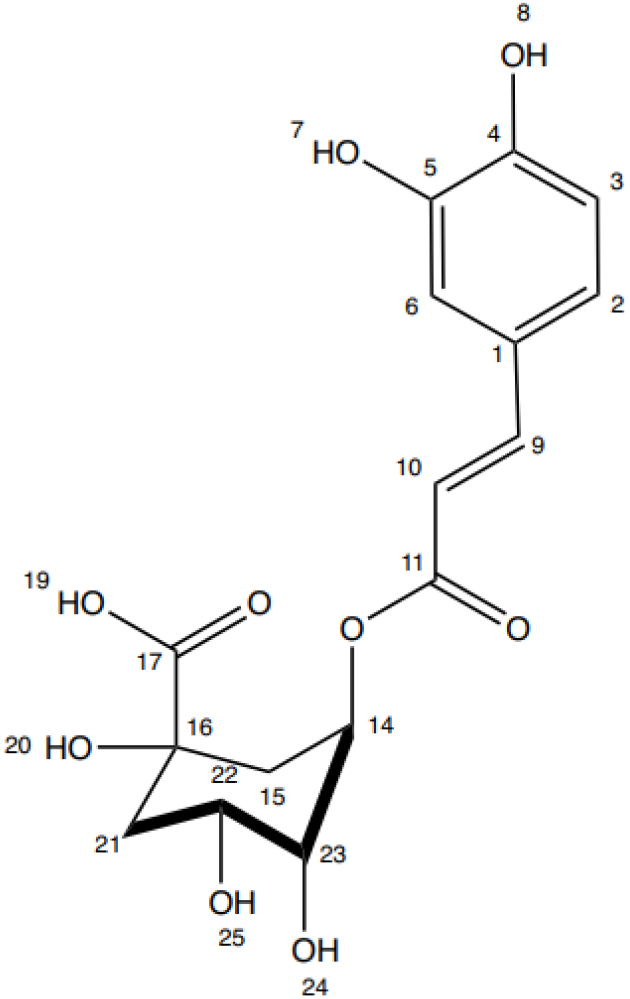
Structural formula of the sample under study, identified as chlorogenic acid.

Signal assignments are presented in Table 3 for the ^1^H and ^13^C{^1^H} spectra.

The data presented in Table 3 confirm that the compound isolated from the red clover callus culture extract fully corresponds, in its chemical structure, to chlorogenic acid. Analysis of the ^1^H NMR spectrum demonstrates the presence of a complete set of proton signals characteristic of the structural fragments of the molecule: caffeic and quinic acids linked by an ester bond. In the downfield region of the spectrum (δ 6.15 – 7.42 ppm), signals of the three aromatic protons of the caffeic acid ring, CH(2), CH(3), and CH(6), are observed as a characteristic ABX system with spin–spin coupling constants of j=2.3 Hz и j=8.8Hz. Signals of the two olefinic protons of the trans double bond, CH(9) and CH(10), are also observed in this region, with a characteristic coupling constant of j=16.6Hz, indicating the trans configuration of this fragment. The aliphatic region (δ 1.81 – 5.55 ppm) contains proton signals of the cyclohexane ring of quinic acid: the diastereotopic methylene groups CH_2_(15) and 1/2CH_2_(21), the oxymethine protons CH(14), CH(22), and CH(23), as well as the hydroxyl groups OH(20), OH(24), and OH(25) of the quinic acid fragment. Signals of the phenolic hydroxyl groups of caffeic acid, OH(7) and OH(8), and of the carboxyl group, OH(19), are observed in the most downfield region (δ 9.18 – 12.45 ppm), which is typical of intramolecularly bonded protons.

**Table 3.**
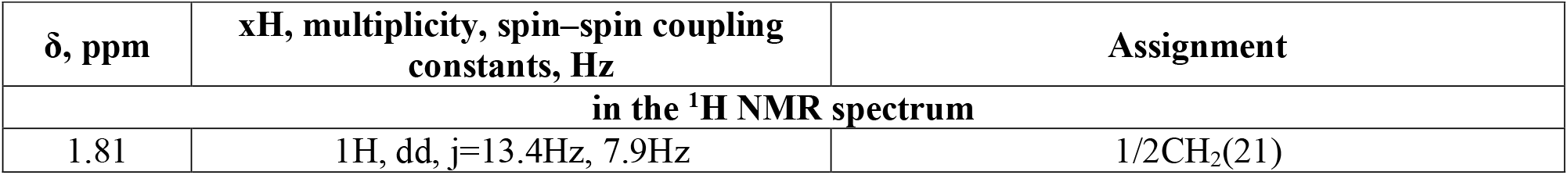
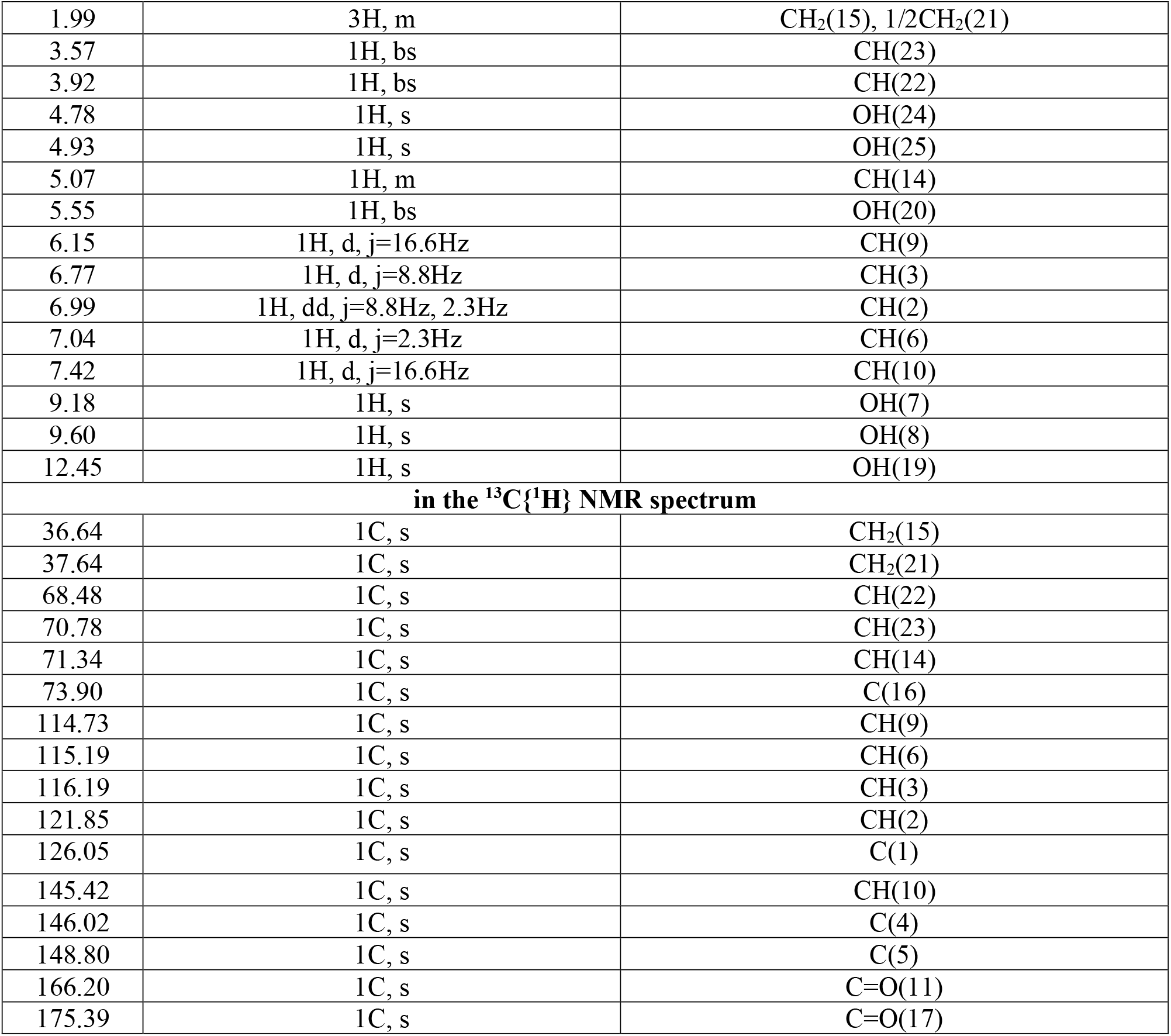
Signal assignments for the chlorogenic acid sample in the ^1^H and ^13^C{1H} NMR spectra.

Signal assignments in the ^13^C{1H} NMR spectrum are fully consistent with the proposed structure. Signals of eight aromatic and olefinic carbon atoms of the caffeic acid fragment were recorded in the range δ 114,73 – 148,80 ppm, including the characteristic signals of the hydroxylated carbon atoms C(4) at δ 146.02 ppm and C(5) at δ 148,80 ppm. Signals of two carbonyl groups were also observed: C=O(11) of the ester bond at δ 166,20 ppm and C=O(17) of the quinic acid carboxyl group at δ 175,39 ppm. The number, multiplicity, and chemical shifts of all recorded signals strictly correspond to the structure of chlorogenic acid. No evidence of impurities was detected in the sample: no extraneous signals were observed in either the proton or carbon spectra. Thus, the combined NMR spectroscopy data indicate the high purity of the isolated compound and complete agreement of its structure with 5-O-caffeoylquinic acid, chlorogenic acid.

## Conclusion

As a result of this study, which employed a set of modern physicochemical methods, such as X-ray diffraction analysis and nuclear magnetic resonance spectroscopy, detailed data were obtained for the first time on the molecular and spatial structure of chlorogenic acid isolated from a 70% ethanol extract of red clover (*Trifolium pratense*) callus culture.

Powder X-ray diffraction showed that the chlorogenic acid sample under study was crystalline, single-phase, and characterized by a high level of purity, consistent with the stated purity of at least 95%. The key crystallographic parameters of the compound were determined: the sample crystallized in an orthorhombic system with unit cell parameters a = 36.7548(5) Å, b = 11.0770(3) Å, c = 7.7947(2) Å, V = 3173.46(11) Å^3^. The high quality of the structural model was confirmed by the low values of the discrepancy factors (R-Bragg = 0.347 %, R_exp_ = 4.75 %, R_wp_ = 5.83 %, R_p_ = 4.39 %) and the goodness-of-fit value (GOF = 1.23 %), indicating the reliability of the results obtained. This diffractogram may serve as a reference pattern for rapid identification and authenticity control of chlorogenic acid.

High-resolution one-dimensional (^1^H and ^13^C{^1^H}) and two-dimensional (^1^H^1^H-COSY, ^1^H^13^C-HSQC, ^1^H^13^C-HMBC) NMR spectroscopy confirmed the chemical structure of the isolated compound, which fully corresponded to 5-O-caffeoylquinic acid, or chlorogenic acid. Complete assignment of all proton and carbon signals in the NMR spectra was performed. The double bond in the caffeic acid fragment was found to have an exclusively trans configuration, as indicated by the characteristic spin–spin coupling constant of j=16.6Hz. Detailed analysis of signal multiplicity and integral intensities, together with the absence of extraneous resonance lines in the ^1^H and ^13^C{^1^H} spectra, confirmed that the sample contained no detectable impurities, which is consistent with the X-ray diffraction data.

Thus, the combined data demonstrate that the biotechnological production scheme used in this study (from *in vitro* callus culture cultivation to isolation and purification of the target product) makes it possible to obtain highly purified chlorogenic acid with a well-defined molecular and crystal structure. The results of this study provide a physicochemical basis for the further standardization of chlorogenic acid as a promising pharmaceutical substance and may be used in the development of regulatory documentation for this product. The set of analytical approaches described here can be recommended as a methodological basis for the structural analysis of other plant-derived polyphenolic compounds.

## Funding

This study was carried out within the framework of the state assignment “Development of biologically active supplements composed of metabolites from in vitro plant objects to protect the population from premature aging” (project FZSR-2024-0008)

The study was performed using the equipment of the Core Facilities “Instrumental Methods of Analysis in Applied Biotechnology” at Kemerovo State University.

